# Conserved gene clusters in the scrambled plastomes of IRLC legumes (Fabaceae: Trifolieae and Fabeae)

**DOI:** 10.1101/040188

**Authors:** Saemundur Sveinsson, Quentin Cronk

**Affiliations:** Faculty of Land and Animal Resources, Agricultural University of Iceland, Keldnaholt, 112 Reykjavik, Iceland; Department of Botany and Biodiversity Research Centre, University of British Columbia, 6270 University Boulevard, Vancouver BC V6T 1Z4, Canada

**Keywords:** Fabeae, IRLC, massively parallel sequencing, plastid operons, plastome evolution, plastome rearrangements, Trifolieae

## Abstract

The plastid genome retains several features from its cyanobacterial-like ancestor, one being the co-transcriptional organization of genes into operon-like structures. Some plastid operons have been identified but undoubtedly many more remain undiscovered. Here we utilize the highly variable plastome structure that exists within certain legumes of the inverted repeat lost clade (IRLC) to find conserved gene clusters. These plastomes exhibit an unusually high frequency of translocations and inversions. We analysed the plastomes of 23 legume species and identified 32 locally collinear blocks (LCBs), which are regions within the plastid genomes that occur in different orientation and/or order among the plastid genomes but are themselves free from internal rearrangements. Several represent gene clusters that have previously been recognized as plastid operons. It appears that the number of LCBs has reached saturation in our data set, suggesting that these LCBs are not random, but likely represent legume plastid operons protected from internal rearrangement by functional constraint. Some of the LCBs we identify, such as *psbD*/C/Z, are previously known plastid operons. Others, such as *rpl32-ndhF-psbA-matK-rbcL-atpB-atpE*, may represent novel polycistronic operons in legumes.

## Introduction

The plastid genome, also known as the plastome, refers to the total genetic information of a single plant organelle, the plastid, which takes many developmental forms, the most notable being the chloroplast (Bock, 2007). Plastid genomes are circular structures of double stranded DNA, usually consisting of about 100-120 genes and are around 120-160 kb long in photosynthesizing plants (Bock, 2007). Their size, structure and gene content are highly conserved across land plants (Wicke *et al.*, 2011). However there are exceptions, such as the Geraniaceae and Campanulaceae, which are two angiosperm families known to contain species with highly rearranged plastomes (Haberle *et al.*, 2008; Guisinger *et al.*, 2011). A dominating feature of plastid genomes is the presence of a large inverted repeat (Wicke *et al.*, 2011; Zhu et al., 2015) separated by a small single copy region that is variable in orientation (Walker et al. 2015). However some plant groups have lost one copy of the repeat, one being a clade within papilionoid legumes (Fabaceae), known as the inverted repeat lost clade (IRLC) (Wojciechowski *et al.*, 2000).

Plants obtained their plastid organelles through an endosymbiosis event with a cyanobacteria-like organism, about 1.5 – 1.6 billion years ago (Margulis, 1970; Hedges *et al.*, 2004). Its bacterial origin gives the plastid genome many prokaryotic features, such as small (70S) ribosomes and the absence of mRNA 3’ polyA tails (see Stern *et al.*, 2010 for a review). An additional ancestral feature of the plastid genome is the organization of its coding region into multiple gene clusters, or operons (Sugita & Sugiura, 1996; Sugiura *et al.*, 1998). These gene clusters are stretches of the plastome consisting of several genes that are transcribed into di-or polycistronic units, which are then processed before translation (Stern *et al.*, 2010). Several such clusters have already been identified in the plastid (Adachi *et al.*, 2012; Ghulam *et al.*, 2013; Stoppel & Meurer, 2013).

Several legume genera within the IRLC are known to harbour highly rearranged plastomes, as a result from multiple translocations and/or inversions: *Trifolium* (Cai *et al.*, 2008; Sabir *et al.*, 2014; Sveinsson & Cronk, 2014), *Pisum* (Palmer & Thompson, 1982), *Lathyrus* (Magee *et al.*, 2010), *Lens* and *Vicia* (Sabir *et al.*, 2014). The aim of this study is to analyse these rearrangements in these genera within IRLC, in order to investigate whether they can be used to study the organization of plastid genomes into operons.

## Material and Methods

### Source of plant material

The plant material for this study came from three sources. First, live plants were collected in the field and transplanted to a glasshouse facility. Secondly, seeds were obtained from a commercial provider, Roger Parsons Sweet Peas (Chichester, UK). Thirdly, seeds were received from the USDA germplasm collection at Pullman, Washington (W6). A full list of germplasm used is given in Table 1. All plants were grown in glasshouse facilities at UBC. In all cases where plants required critical determination they were grown until flowering, and herbarium voucher specimens were then collected (UBC).

**Table 1.** Information regarding the plastid genomes used in this study. Details regarding the Illumina sequencing and voucher details are presented where applicable. An asterisk (*) indicates species sequenced in this study.

| **Species**<br>**(USDA seed accession)** | **GenBank**<br>**accession** | **Herbarium**<br>**voucher (UBC<sup>2</sup>)** |
| --- | --- | --- |
| *Cicer arietinum* L. (NA) | NC_011163 | NA |
| *Medicago truncatula* Gaertn.<br>(NA) | NC_003119 | NA |
| * *Trifolium strictum* L.<br>(PI 369147) | KJ788292 | V241491 |
| *T. grandiflorum* Schreb. (NA) | NC_024034 | NA |
| *T. aureum* Pollich (NA) | NC_024035 | NA |
| * *T. boissieri* Guss. (PI 369022) | KJ788284 | V241490 |
| * *T. glanduliferum* Boiss. (PI<br>296666) | KJ788285 | V241492 |
| * *Vicia sativa* L. (PI 293436) | KJ850242 | NA |
| * *Lens culinaris* Medik.<br>(PI 592998) | KJ850239 | NA |
| * *Pisum sativum* L. (W6 32866) | KJ806203 | V241498 |
| * *Lathyrus clymenum* L. (RP<sup>3</sup>) | KJ850235 | V241501 |
| * *L. tingitanus* L. (RP<sup>3</sup>) | KJ850238 | V241502 |
| *L. sativus* L. (NA) | NC_014063 | NA |
| * *L. odoratus* L. (RP<sup>3</sup>) | KJ850237 | V241503 |
| * *L. pubescens* Hook. & Arn.<br>(RP<sup>3</sup>) | KJ806200 | V241505 |
| * *L. inconspicuus* L. (W6 2817) | KJ850236 | V241504 |
| * *L. davidii* Hance (RP<sup>3</sup>) | KJ806192 | V241506 |
| * *L. palustris* L. (NA) | KJ806198 | V241511 |
| * *L. japonicus* Willd.(NA) | KJ806194 | NA |
| * *L. littoralis* Endl.(NA) | KJ806197 | NA |
| * *L. graminifolius* (S.Watson)<br>T.G.White (DLP<sup>4</sup> accession:<br>920239) | KJ806193 | V241507 |
| * *L. ochroleucus* Hook.<br>(NA) | KJ806199 | V241489 |
| * *L. venosus* Muhl. ex Willd.<br>(NA) | KJ806202 | V241509 |

<sup>1</sup>SD: Standard Deviation
<sup>2</sup>UBC Herbarium, Vancouver BC Canada
<sup>3</sup>Roger Parsons Sweet Peas, Chichester UK
<sup>4</sup>Desert Legume Project Germplasm, AZ USA

### Illumina sequencing

Total DNA was extracted from fresh leaf material following a modified version of the CTAB protocol (Doyle & Doyle, 1987). RNase treatments were performed (cat. 19101, QIAGEN, Germantown, MD) and DNA quality was assessed by visual inspections on 1% agarose gels. Illumina sequencing libraries were constructed from high quality DNA, using the NEXTflexTM DNA sequencing kit (100 bp Paired-End reads) (cat: 5140-02, BiooScientific Corp, TX). We followed the manufacturer’s protocol and c. 400 bp DNA fragments were size selected using Agencourt AMPure XpTM magnetic beads (cat. A63880, BeckmanCoulter Genomics, MA). Completed libraries were pooled and sequenced on a lane of the Illumina HiSeq-2000 platform.

### Plastid genome assemblies and annotation

Trimmomatic v.0.3 (Lohse *et al.*, 2012) was used to trim and remove low quality Illumina reads, with the following flags: LEADING:20 TRAILING:20

SLIDINGWINDOW:4:15MINLEN:36. High quality reads were used in all subsequent analysis and singlet reads, i.e. reads without a paired end, were discarded. We used the *de novo* method implemented in CLC Genomic Workbench v.7.0.2 to generate assemblies for each species, using the default settings. Contigs of plastid origin were identified by a BLASTN search (Altschul *et al.*, 1997) to a plastid genome of a closely related species. These were generally the largest and most high coverage contigs in the *de novo* assembly and always had an E-value of 0 when blasted to the reference plastome. Regions with nucleotides scored as Ns were manually resolved by retrieving sequence information directly from the quality-trimmed reads. For most species, the *de novo* assembly returned a single large plastid contig. When needed, multiple contigs containing plastid sequence were joined by hand, using information from the quality-trimmed reads. The quality of each plastome assembly was verified by visually by inspecting a BWA mem pileup, v. 0.7.5a (Li & Durbin, 2009), of paired end reads using Tablet v.1.13.12.17 (Milne *et al.*, 2013). We made sure that the connections between manually joined contigs were supported by paired-end read mapping. Finally all plastome assemblies were annotated using DOGMA (Wyman *et al.*, 2004).

### Phylogenetic analysis

Due to the extensive rearrangements observed in the plastomes (see Sabir *et al.*, 2014; Sveinsson & Cronk, 2014), we restricted our plastome phylogenetic analysis to protein coding genes. We used a custom phylogenetic pipeline, plast2phy, that extracted protein coding regions from DOGMA annotated plastomes, aligned individual gene with Mafft v. 7.0.5(-auto flag) (Katoh & Standley, 2013), trimmed alignment gaps using trimAl v.1.2 (-automated1 flag) (Capella-Gutiérrez *et al.*, 2009) and finally generated a concatenated alignment of all genes. The pipeline, Plast2phy, written in Python, is available at https://github.com/saemi/plast2phy. Model of base substitution were tested for the concatenated matrix using jModelTest v.2.1.1 (Guindon & Gascuel, 2003; Darriba *et al.*, 2012). Using the Akaike information criterion (AIC), we determined the GTR+G+I model optimal for the concatenated plastome alignment. We analysed the dataset under maximum likelihood (ML; Felsenstein, 1973) using GARLI (Zwickl, 2006). We ran GARLI v. 2.0 with default settings, using ten independent searches and 100 bootstrap replicates. Bootstrap consensus was calculated using SumTrees v. 3.3.1 in the DendroPy package (Sukumaran & Holder, 2010). Trees from phylogenetic analysis were drawn using FigTree v.1.4.0 (http://tree.bio.ed.ac.uk/software/figtree/), rooted with *Cicer aretinum*.

### Identification of locally collinear blocks (LCBs) in plastid genomes

The progressive alignment method, implemented in the MAUVE v.2.3.1 package (Darling *et al.*, 2010), was used with the default parameters to identify locally collinear blocks (LCBs) among the plastid genomes listed in Table 1. In this study, a LCB represents a region within a plastid genome that can occur in different orientation and/or order among the studied plastid genomes, but is free from any internal rearrangements (see Darling *et al.*, 2010). These regions are therefore putatively orthologous in nature. I used two programs, projectAndStrip and makeBadgerMatrix (downloaded from http://gel.ahabs.wisc.edu/mauve/snapshots/, on 11 November 2014) to generate a LCB boundary file from the MAUVE alignment. The LCB boundary file contained information on where the LCBs start and end in each of the analysed plastome. I used the chickpea plastome (*Cicer arietinum)* [NCBI Reference Sequence NC_011163] (Jansen *et al.*, 2008), as the reference plastome. We visualized the observed plastid rearrangements using Circos v.0.66 (Krzywinski *et al.*, 2009), were a custom Python script was to generate Circos input files and executing the program for each of plastid genome (Figs. 1 and 2). Figure 2 was generated by manually combining all the Circos maps with a phylogenetic cladogram using inkscape (www.inkscape.org). Information regarding the size of LCBs and lengths of protein coding genes in Table 2 are based on the reference plastome of *Cicer arietinum* [NC_011163]. Putative unannotated tRNAs were identified using MITOS (Bernt *et al.*, 2013).

**Table 2.**
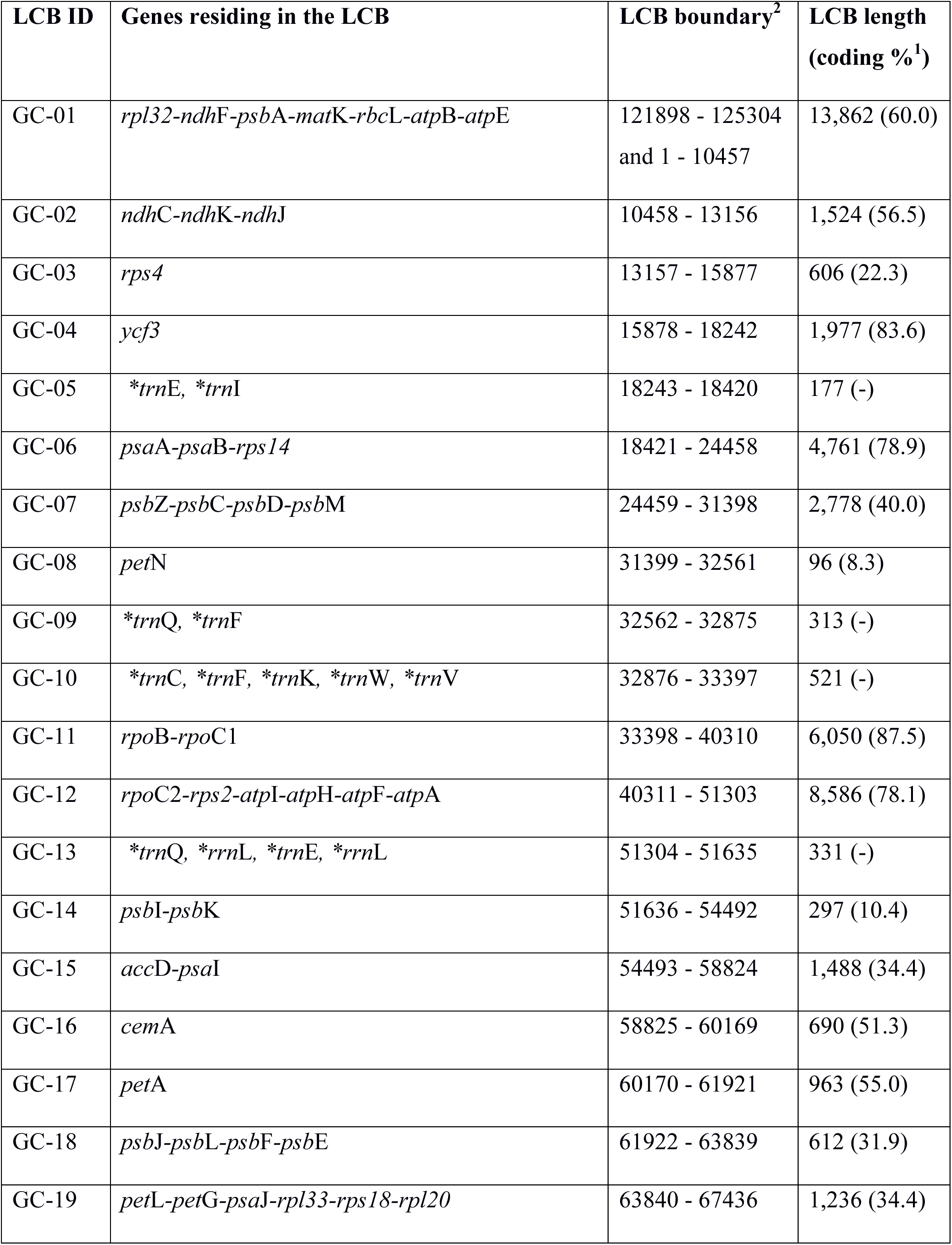
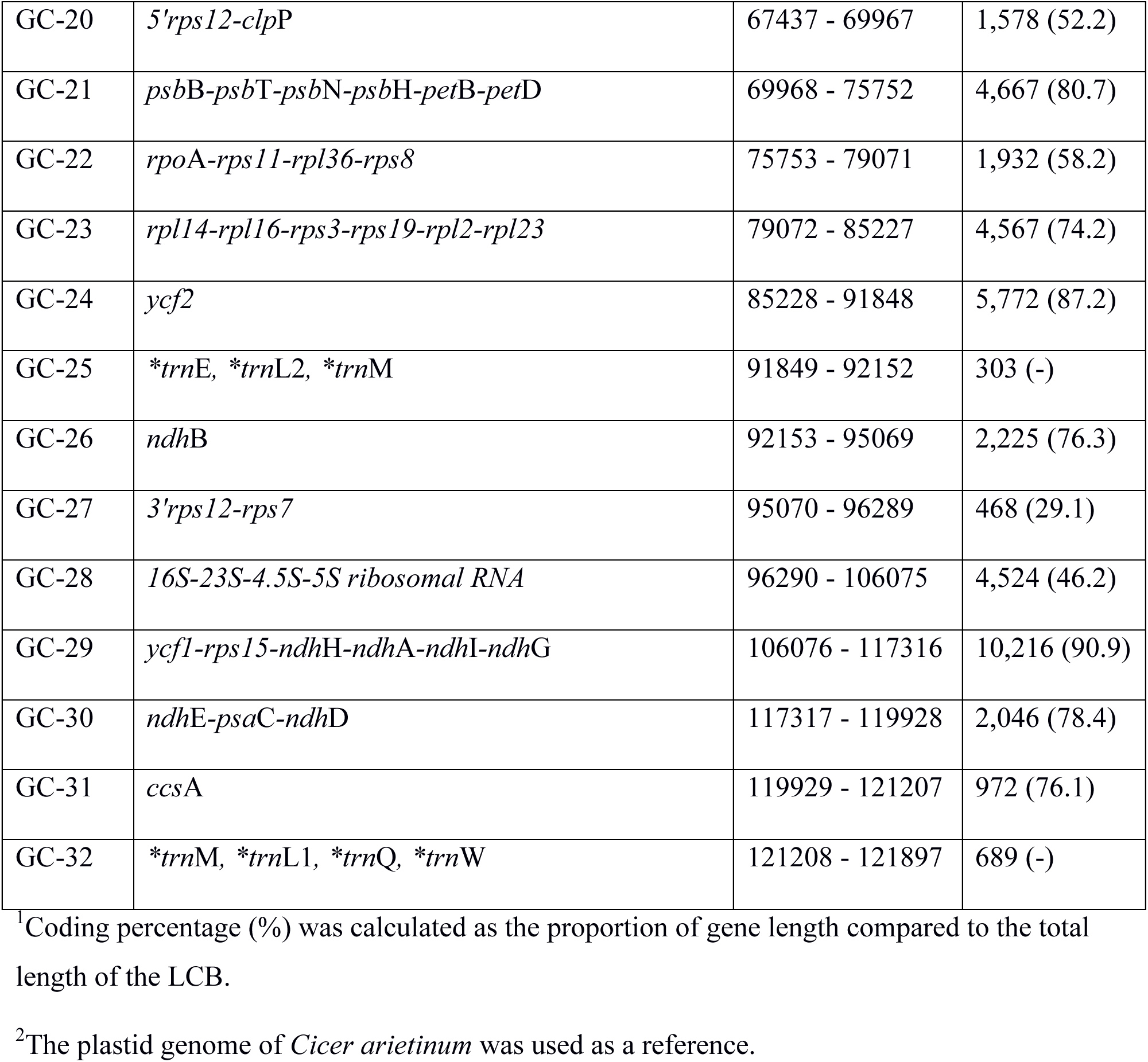
Details regarding the gene content, positional boundaries, length of locally collinear blocks identified within the analysed plastome. An asterisk (*) marks putative tRNAs unannotated in the *Cicer* plastome.

**Figure 1.**
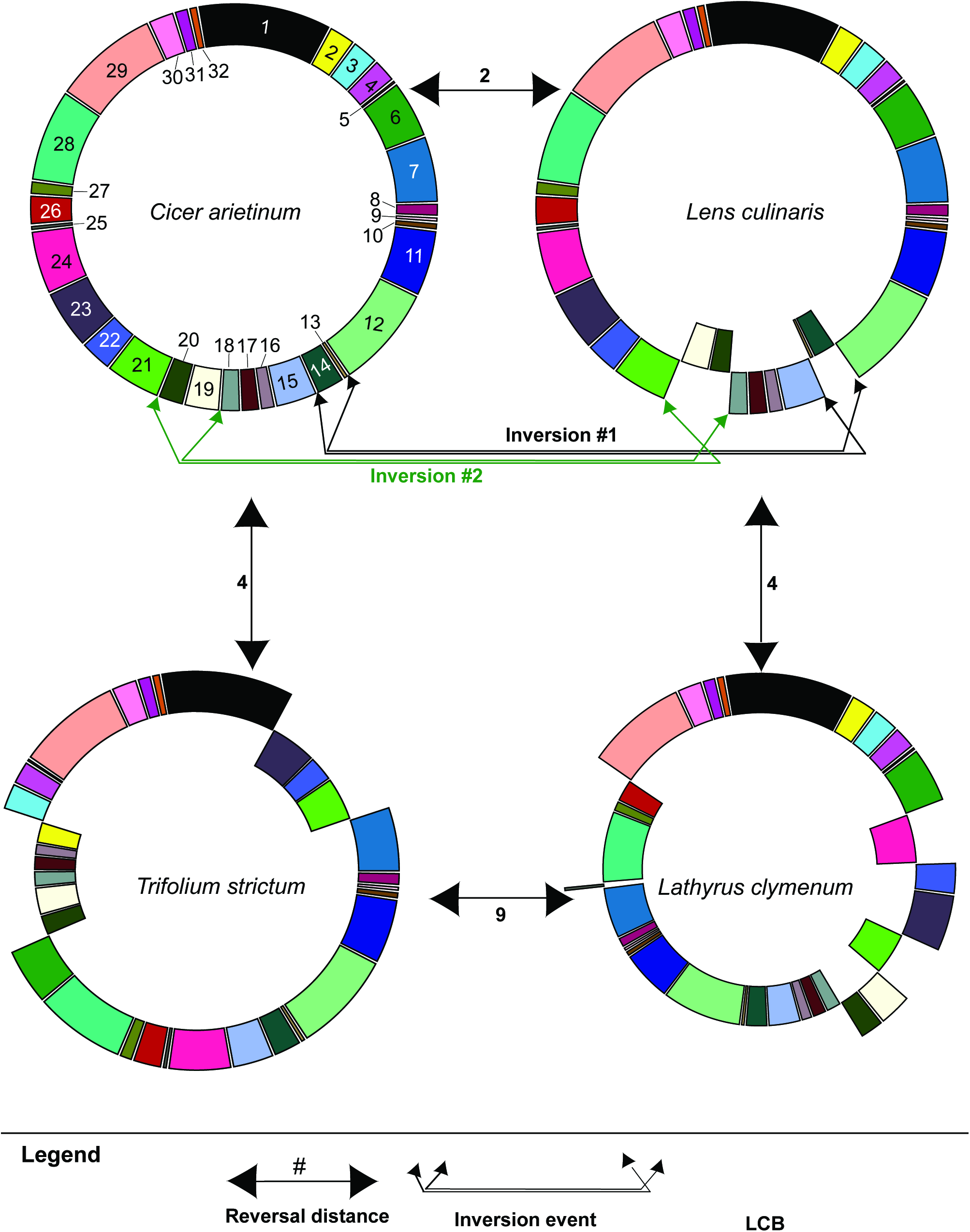
Example of rearrangements and reversal distances between plastid genomes. A comparison of the order and orientation of locally collinear blocks (LCBs) among four species: *Cicer arietinum, Lens culinaris, Trifolium strictum* and *Lathyrus clymenum*. LCBs are represented as differently coloured boxes and the numbers refer to the LCB IDs in Table 2. Inverted LCBs are positioned on an inner circle. Reversal distances (see Material and Methods) between species is shown on arrowed lines. Two plastid inversion events between *C. arietinum* and *L. culinaris* are highlighted.

**Figure 2.**
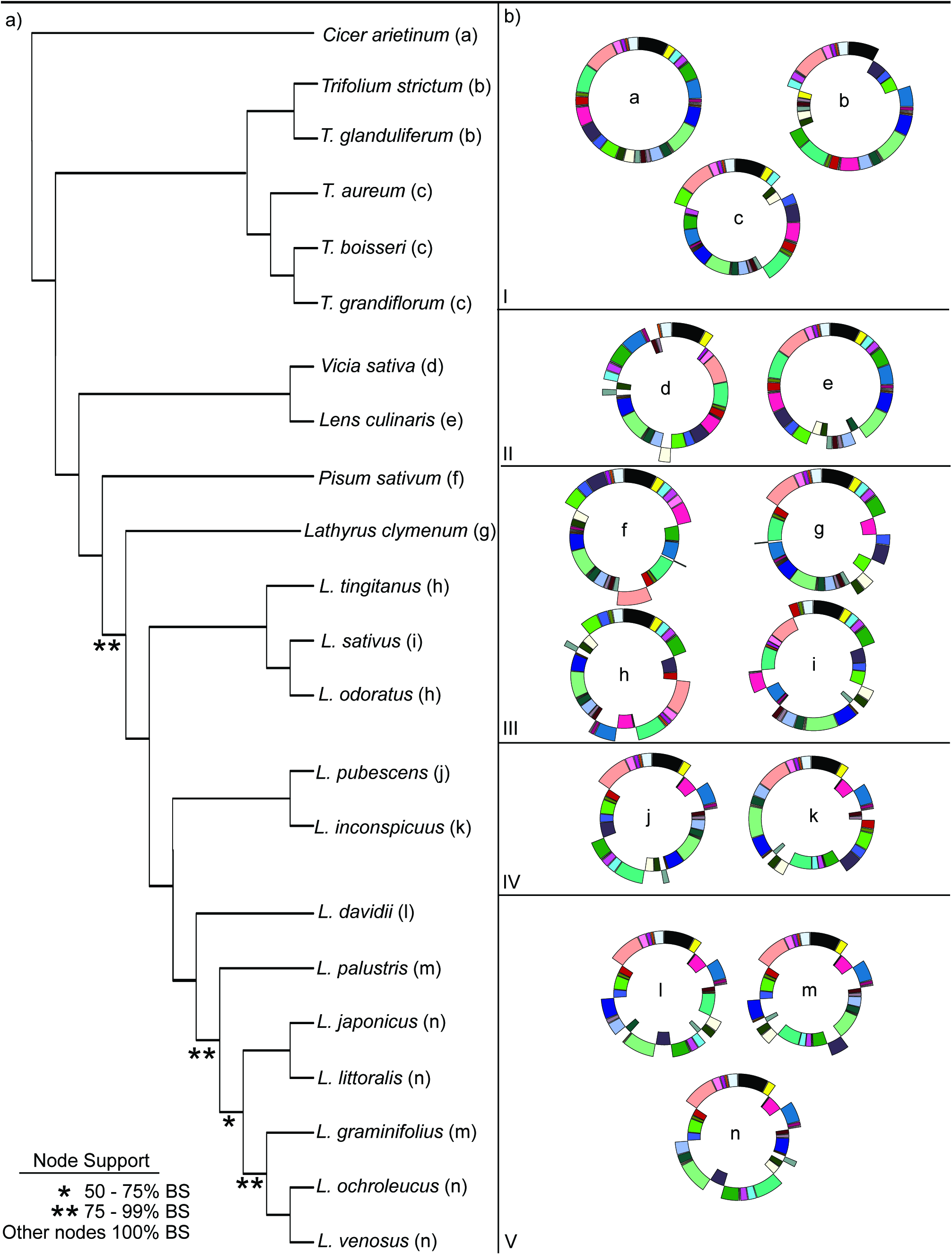
Plastid rearrangements of the investigated species shown in a phylogenetic context. (a) A cladogram showing the phylogenetic relationship among the species in this study. The phylogeny was constructed from a concatenated alignment of plastid protein coding genes. All nodes except one were retrieved with a 100% bootstrap support. (b) Visual representation of the rearranged plastid genomes of the species in this study. Locally collinear blocks (LCBs) are represented as coloured boxes. The letter in the middle of each circle refers to the major plastotype of each species, shown after the species names in parenthesis in the cladogram (a).

### Relationship among divergence time and plastid rearrangements

In order to investigate the relationship among species divergence and the number plastid rearrangements, we estimated two relevant parameters in a pairwise manner. Firstly we estimated the reversal distance using GRIMM v. 2.0.1 (Tesler, 2003). Reversal distance is the minimum number of reversal steps for two genomes to become completely syntenic (Tesler, 2003). GRIMM uses the LCB boundary file, described in the previous section, as its input file. Secondly we used the synonymous substitution rate (Ks) between pairs of plastid genomes as an indicator for the divergence between species. Ks values were calculated using MEGA v.6.0 (Tamura *et al.*, 2013), from a concatenated alignment file of the plastid protein genes. Divergence times can be estimated using published estimates of plastid mutation rates, which range from 1.1 – 2.9 silent substitutions per billion years (Wolfe *et al.*, 1987). The reversal distance was plotted against divergence time using R v. 3.0.2 (R Core Team, 2014) and ggplot2 (http://ggplot2.org/), and their relationship visualized using a smoothing curve, using the following command: geom_smooth(degree=1, shape=2/3, method=‘loess’, level=.95).

## Results

### Phylogeny of species with rearranged plastomes

Mapping the plastome architecture onto the phylogenetic tree of the investigated legume species indicates that the plastomes have clearly undergone multiple multiple and rounds of inversions and translocations throughout the tree (Figs. 1, 2a and 2b.) The phylogenetic analysis of the protein coding regions of our completely sequenced plastomes proved useful in resolving the relationships among the studied species (Fig. 2a.). The phylogeny reported here largely confirms previous studies (LPWG 2013) and is consistent with the *Trifolium* phylogeny using the same methods reported previously (Sveinsson & Cronk 2014) and Fabeae (previously reported in Magee *et al.*, 2010). The tree is rooted on *Cicer*, which along with *Medicago*, shows no evidence of plastid rearrangements compared to other IRLC legumes (Supporting Information 1) and, with the exception of the lost inverted repeat (Wojciechowski *et al.*, 2000), their plastomes are collinear with *Lotus japonicus* (Supporting Information 1, Fig. 3).

**Figure 3.**
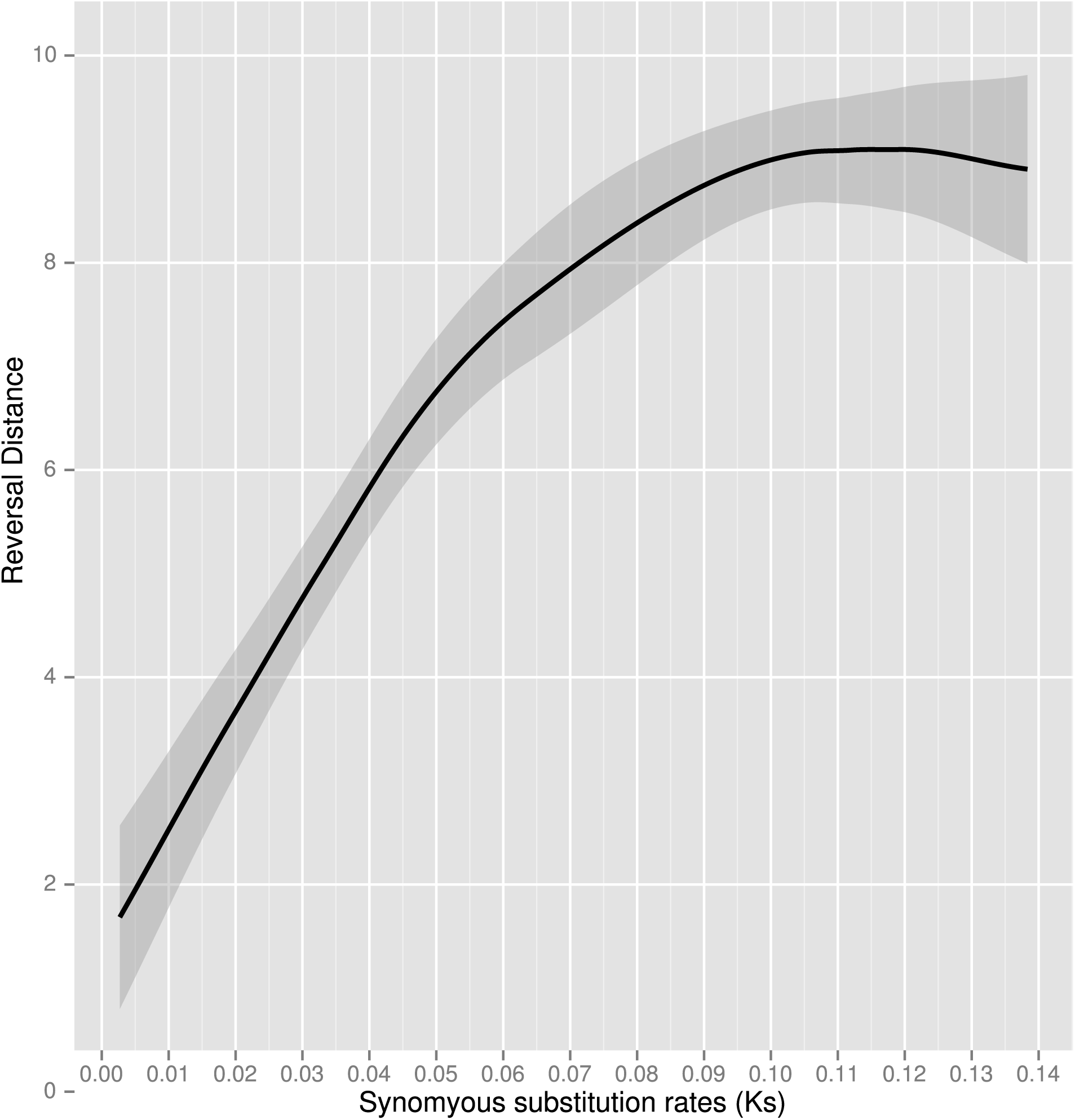
The relationship between the pairwise species divergence and the reversal distance of the plastid genome analysed in this study. Synomyous substitutions rates (Ks) of plastid protein coding genes are used as a proxy for species divergence time, and are shown on the x-axis. The reversal distance (extent of rearrangements) is shown on the y-axis. The black line is a smooth curve that illustrates the relationships among these two variables. 95% confidence intervals are represented as grey areas around the line.

### Many locally collinear blocks (LCBs) correspond to previously reported plastid operons

The entire plastid genome in these species has been broken up multiple times by rearrangement but certain genomic blocks have never been broken up. MAUVE identified a total of 32 localized collinear blocks (LCB) (Fig. 2b) in the 23 analysed plastomes. Out of these 32 LCBs, 26 contained protein-coding genes and one LCB was made up of the plastid rRNA genes (see Table 2). These 26 blocks varied in size and contained gene clusters (GCs) that varied in the number of genes that they encompass (see Table 2). Nine of the blocks (gene clusters) contained only a single gene, five blocks were composed of two genes and the remaining 12 blocks consisted of more than 2 genes. The largest gene cluster (GC) is GC-1, 13.8 kb in length, containing the following genes: *rpl32, ndhF, psbA, matK, rbcL, atpE* and *atpB* (Table 2). The smallest gene cluster detected was GC-8, about 1.2 kb in length, containing only a single gene, *petN.* Many of the gene clusters have previously been recognized as plastid operons (i.e. transcriptional units), such as GC-2, 6, 7, 18, 27 and 31 (see Sugita & Sugiura, 1996). Several other gene clusters share extensive similarities with previously reported plastid operons but can differ in the presence or absence of a single gene, e.g. GC-11, 12 and 21. Gene containing LCBs cover about 98% of the total length of the *Cicer arietinum* plastome. This suggests that the delimitation of these clusters is not random and is under functional constraint (see discussion).

### The number of plastome rearrangements increases with divergence time, but levels off

We investigated the relationship between species divergence and the number of plastome rearrangements, by plotting sequence evolution against genome rearrangement. Specifically we plotted pairwise synonymous substitution rate (Ks) against pairwise reversal distances (see Fig. 3). The rationale for this analysis was that if formation of new LCBs in the plastomes is constrained by the presence of plastid operons, blocks should increase in number until functional constraint does not allow further break-up of blocks. We find a pattern consistent with this constraint hypothesis. With evolutionary distance (approximating time) rearrangements increase until saturation is reached. When a smoothing curve is fitted through the data, we observe what seems to a strong positive correlation between evolutionary divergence and plastome rearrangements up until about Ks~0.10 where it starts to level off, which relates to about 9 in reversal distance (Fig. 3). If the plastomes were under no functional constraint there would be no obvious reason that the relationship between divergence and reversal distance would level off at that point. Our results suggest that there is functional constraint on the observed plastome rearrangements and its most likely source is the preservation of functional di-and polycistronic plastid transcriptional units (see discussion). This functional constraint appears to place a limit on the number LCBs (Fig. 3), i.e. limit the extent to which blocks of genes can be broken up. The block cannot be further divided, by inversion and translocation, without breaking co-transcriptional units.

## Discussion

### The rearrangements of plastomes in IRLC legumes

Plastid genomes of analysed *Trifolium* and the Fabeae (*Lens, Vicia, Pisum* and *Lathyrus*) species are highly rearranged, as a result of multiple rounds of translocations and/or inversions (Fig. 2 and Fig. 2). These rearrangements have previously been reported (Palmer & Thompson, 1982; Cai *et al.*, 2008; Magee *et al.*, 2010; Sabir *et al.*, 2014; Sveinsson & Cronk, 2014). Plastid genomes tend to be quite conserved structurally across land plants (see Wicke *et al.*, 2011). However, besides these IRLC legumes, there are other well-known exceptions, such as Geraniaceae (Guisinger *et al.*, 2011) and Campanulaceae (Haberle *et al.*, 2008). The plastome rearrangements described here for the Fabeae appear to be most similar to those reported in *Trachelium caeruleum* (Campanulaceae), since they do not involve proliferation of repeated elements, such as in certain *Trifolium* species (Sveinsson & Cronk, 2014) or in the Geraniaceae (Guisinger *et al.*, 2011; Weng *et al.*, 2013). The functional cause of these rearrangements is not known. The stability of plastid genomes is maintained through recombinational mechanisms, which are controlled by a large number of nuclear genes (see Maréchal & Brisson, 2010). The loss of the inverted repeat may be involved in the genome instability but, if so, the relationship is not simple as some IRLC legumes such as *Medicago* and *Cicer* have well conserved typical legume gene orders. Whatever the functional changes that result from this unprecedented genome instability, it is clear that it offers a unique opportunity for the study the organization of inviolable transcriptional units within the plastid genomes of flowering plants. Against a background of extensive genome scrambling, blocks with conserved gene order stand out.

### Do conserved blocks in otherwise rearranged plastomes represent operons?

The genespace of plastid genomes is organized into transcriptional units, similar to operons in the genome of their cyanobacterial ancestors (Stern *et al.*, 2010). However it is important to note that despite plastids being of bacterial origin, most aspects of the regulation of plastid gene expression are radically different from bacteria, mainly due to interactions with the nuclear genome (Stern *et al.*, 2010). Nevertheless, it is well established from functional studies that many plastid genes are organized into dicistronic or polycistronic operon-like units, i.e. co-regulated gene blocks, also known as transcriptional units (Sugita & Sugiura, 1996). It is therefore reasonable to assume that any structural rearrangements that would break up these transcriptional units would be detrimental to the plastid and be selected against.

Our results are in agreement with that assumption, as many of the gene clusters that we observe are known plastid polycistronic operons (Table 2 here; and Table 2 in Sugita & Sugiura, 1996). Examples of this are: (i) Gene Cluster 21 (GC-21, Table 2) that seems to correspond to the *psbB* operon, which has been extensively studied (Stoppel & Meurer, 2013); (ii) GC-7 which contains the same genes as the *psbD*/C/Z operon, which has been characterized in tobacco (Adachi *et al.*, 2012); (iii) GC-28 which contains all the rRNA genes, which are necessary to construct the plastid 70S ribosome. The numerous genes that are not associated with any other and freely translocate independently, are likely to represent single gene transcriptional units, i.e. monocistronic operons. Gene Cluster 1 (*rpl32-ndhF-psbA-matK-rbcL-atpB-atpE*) is of particular interest, as it contains seven cistrons that previously were thought to be transcribed independently (i.e. as monocistronic units) or belong to different operons (see Table 2 in Sugita & Sugiura, 1996). Our results are highly suggestive that that GC-1 is a conserved plastid operon, at least in the legume species analysed here. Six LCBs without any annotated protein-coding or RNA genes were also identified (varying in size between 177 and 689 nt (see Table 2; Supporting Information 2). However, we identified putative unannotated tRNAs in these blocks (Table 2) and so it is possible that they too are under functional constraint.

These results demonstrate that identification of conserved gene clusters in this clade of rapid structural evolution is a powerful way of provide evidence for previously described plastid operons and potentially to find new ones. Such is the extent of the genic reorganization in the sampled species that it may be argued that the persistence of multiple intact gene blocks is implausible unless these units (Table 2) are inviolable as they represent the fundamental regulatory architecture of the legume plastid.

**Table 3.**
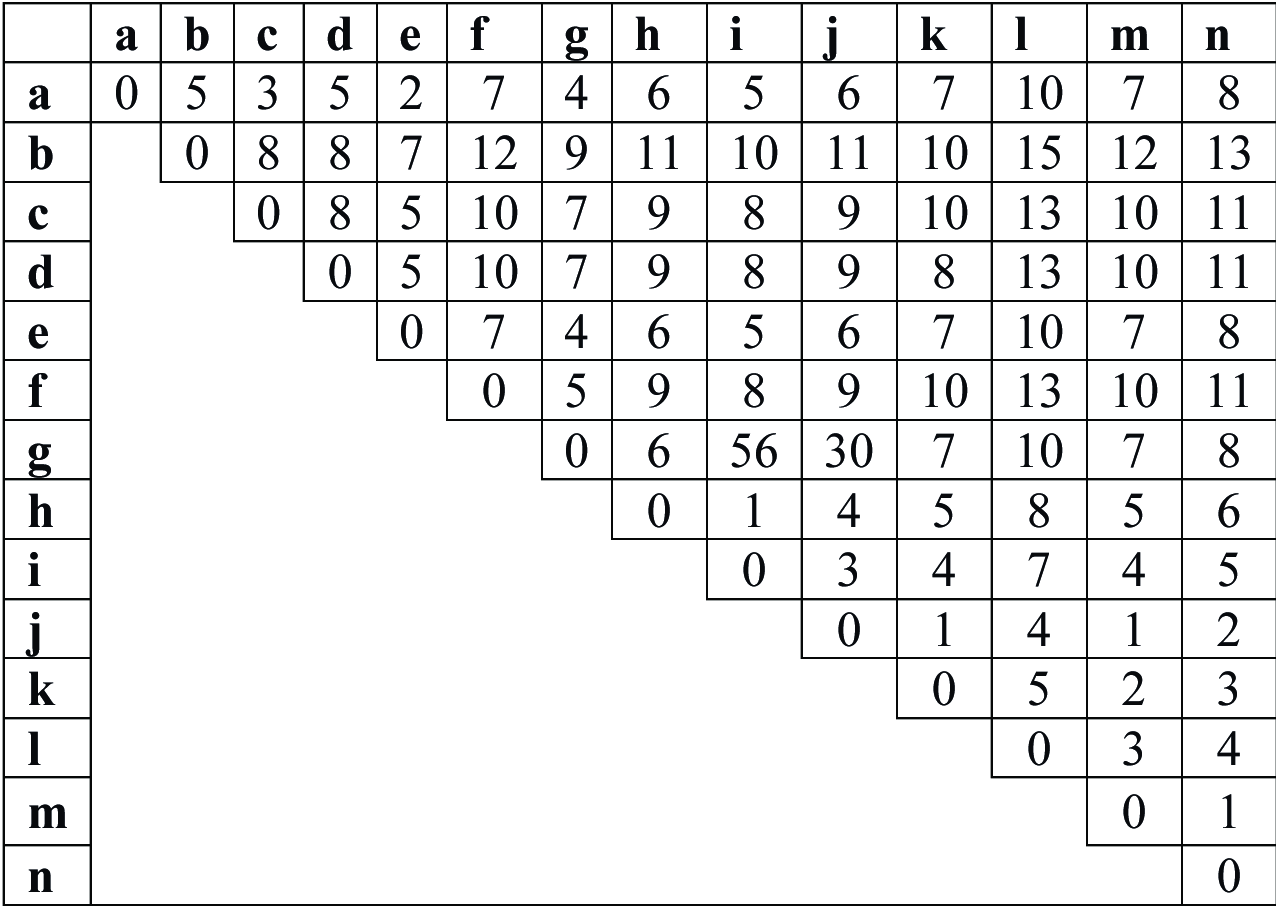
Reversal distance matrix, produced by GRIMM, between each major plastotypes (see Fig. 2b).

|  | **a** | **b** | **c** | **d** | **e** | **f** | **g** | **h** | **i** | **j** | **k** | **l** | **m** | **n** |
| --- | --- | --- | --- | --- | --- | --- | --- | --- | --- | --- | --- | --- | --- | --- |
| **a** | 0 | 5 | 3 | 5 | 2 | 7 | 4 | 6 | 5 | 6 | 7 | 10 | 7 | 8 |
| **b** |  | 0 | 8 | 8 | 7 | 12 | 9 | 11 | 10 | 11 | 10 | 15 | 12 | 13 |
| **c** |  |  | 0 | 8 | 5 | 10 | 7 | 9 | 8 | 9 | 10 | 13 | 10 | 11 |
| **d** |  |  |  | 0 | 5 | 10 | 7 | 9 | 8 | 9 | 8 | 13 | 10 | 11 |
| **e** |  |  |  |  | 0 | 7 | 4 | 6 | 5 | 6 | 7 | 10 | 7 | 8 |
| **f** |  |  |  |  |  | 0 | 5 | 9 | 8 | 9 | 10 | 13 | 10 | 11 |
| **g** |  |  |  |  |  |  | 0 | 6 | 56 | 30 | 7 | 10 | 7 | 8 |
| **h** |  |  |  |  |  |  |  | 0 | 1 | 4 | 5 | 8 | 5 | 6 |
| **i** |  |  |  |  |  |  |  |  | 0 | 3 | 4 | 7 | 4 | 5 |
| **j** |  |  |  |  |  |  |  |  |  | 0 | 1 | 4 | 1 | 2 |
| **k** |  |  |  |  |  |  |  |  |  |  | 0 | 5 | 2 | 3 |
| **l** |  |  |  |  |  |  |  |  |  |  |  | 0 | 3 | 4 |
| **m** |  |  |  |  |  |  |  |  |  |  |  |  | 0 | 1 |
| **n** |  |  |  |  |  |  |  |  |  |  |  |  |  | 0 |

## Acknowledgements

We thank C. Hefer with computer help, D. Kaplan for greenhouse assistance, the Western Canada Research Grid (Westgrid) for computer facilities, USDA and the Desert Legume project for seeds, A. Kuzmin for assistance with sequencing library prep. The work was funded by the Natural Sciences and Engineering Research Council of Canada (NSERC) Discovery Grants Program (grant no. RGPIN-2014-05820 grant to QC).

